# Do place cells dream of deceptive moves in a signaling game?

**DOI:** 10.1101/2022.01.30.478398

**Authors:** André A. Fenton, José R. Hurtado, Jantine A.C. Broek, EunHye Park, Bud Mishra

## Abstract

We consider the possibility of applying game theory to analysis and modeling of neurobiological systems. Specifically, the basic properties and features of information asymmetric signaling games are considered and discussed as having potential to explain diverse neurobiological phenomena at levels of biological function that include gene regulation, molecular and biochemical signaling, cellular and metabolic function, as well as the neuronal action potential discharge that can represent cognitive variables such as memory and purposeful behavior. We begin by arguing that there is a pressing need for conceptual frameworks that can permit analysis and integration of information and explanations across the many scales of diverse levels of biological function. Developing such integrative frameworks is crucial if we are to understand cognitive functions like learning, memory, and perception. The present work focuses on systems level neuroscience organized around the connected brain regions of the entorhinal cortex and hippocampus. These areas are intensely studied in rodent subjects as model neuronal systems that undergo activity-dependent synaptic plasticity to form and represent memories and spatial knowledge used for purposeful navigation. Examples of cognition-related spatial information in the observed neuronal discharge of hippocampal place cell populations and medial entorhinal head-direction cell populations are used to illustrate possible challenges to information maximization concepts. It may be natural to explain these observations using the ideas and features of information asymmetric signaling games.

## INTRODUCTION

A child might ask “how does the brain work?” Because computers are now second nature and the brain clearly computes, one might be tempted to draw on analogies between brains and computers to answer the child, but an agile self-respecting neuroscientist is more likely to answer, “well, you know, the brain is the most complex system we know of, so it should not surprise you that we do not understand how it works.” Dinner party guests often ask “how does memory work?” to which our neuroscientist might answer, “your neurons make a molecule named PKMzeta that is crucial,” or “synapses increase their effectiveness to store memories,” or “when you experience something, the neurons in your brain discharge electrical signals in specific patterns, those patterns replay when you recall the memory,” or “there is a part of the brain called the hippocampus, it is where you store memories, until they are transferred to the neocortex.” Depending on the quality of the wine, the neuroscientist might add “it doesn’t seem at all like computer memory.” While each of these proximate explanations is in a sense standard, it is remarkable that each involves a distinct level of biological organization, where a great amount of self-consistent, rigorously-obtained detail is known within the level, but rather little is known about how to connect the phenomena between the levels and move forward to an ultimate explanation. Such an explanation would also encompass evolution, development, and genetics. Indeed, concepts like transcription, translation, and post-translation modifications like phosphorylation that operate at the nanoscale and minutes-long timescales of genes and macromolecules, may appear off the mark when explanations turn to the dynamics of electrical discharge through neuronal circuits and their computations. It doesn’t seem at all like computer memory in terms of implementation, but what about in terms of the algorithms or computations that define memory?

The experimentalists amongst us, work to understand how the hippocampus-entorhinal cortex neuronal circuitry operates. We identified that the persistent, autophosphorylating kinase, PKMzeta (PKMζ) is translated from mRNA at postsynaptic dendritic sites that were activated during recent learning and that this metabolic synthesis is crucial for long-term memories to persist (Sacktor, 2011;Tsokas et al., 2016). We identified that increased PKMζ expression and synaptic strengthening persists at a subset of hippocampal-entorhinal synapses, for at least a month, so long as the memory persists (Pavlowsky et al., 2017;Hsieh et al., 2021). We also validated the hypothesis that memories of a place in which discomfort was experienced, are recollected when slow gamma oscillations originating at the Schaffer collateral synapses of hippocampus subfield CA1 dominate mid-frequency gamma oscillations that originate at the *stratum lacunosum moleculare*. This competition manifests as memory-associated neuronal ensemble “place cell” discharge that resembles the ensemble location-specific discharge at the recollected place, despite the subject initiating the recollection from the current location, which is a physically different place (Dvorak et al., 2018;Dvorak et al., 2021). Although we study these memory phenomena at distinct levels of biology, in the same laboratory, we still shy away from designing experiments to test hypotheses across the different levels of biology. This is in large part because we lack conceptual frameworks that are useful for describing, organizing, or understanding the cross-level neurobiological memory phenomena (see Bell, 2008).

## THE CHALLENGE OF CROSS-SCALE ANALYSIS AND UNDERSTANDING

The increasing computational abilities of computers have certainly inspired conceptual bridge-building between neuronal computations operating in brains and the computations within artificial neural networks that occur within computers. A key conceptual distinction is that unlike typical computers, brains are self-organizing and changing. Simply using a brain changes it structurally as well as functionally, so that it will operate differently in the future, as we recently showed in mice (Chung et al., 2021). Among the many other less conceptual differences, a key practical distinction between brains and computers is that we have complete information on the latter’s processing, a ground truth of sorts because we know the precise mechanisms that define a microprocessor’s structure and function – in short, using the Marr framework for memory (Marr, 1971), we know the implementation, algorithms, and the computations, completely. In contrast, with regards to the brain, we have incomplete knowledge of the myriad neurobiological processes that result in neuronal implementations, algorithms, and even the computations. We believe this severely limits the generalizability of analogies between the two computing systems; the analogies are especially severely strained by attempts to translate concepts across the different scales of neurobiological function that brains operate within (Jonas and Kording, 2017). Nonetheless, analyzing the brain as an information processing organ is both commonplace and has been especially generative for systems neuroscience, motivated by techniques that are germane to computer science like information theory, circuit analysis, and computational simulations.

Despite the dramatic increase in our ability to record the neuronal discharge from larger and larger ensembles of neurons from freely-behaving subjects, the unavoidable under-sampling of neurons has motivated an interest in a theoretical framework that can guide predictive models from sparse data with the intention to unify observations at many scales of analysis (Bassett and Sporns, 2017). It is here where a new analogy may have substantial utility if it can explain unique biochemical interactions that are not found in computers, and also account for the influence and merit of brain-computer analogies for systems neuroscience.

We believe that the concepts and analytical framework of signaling games is well suited for this cross-scale analysis. The framework has been applied to diverse domains of enquiry, where information is asymmetrically distributed among freely interacting entities. Applications have included the evolution of the genetic code (Jee et al., 2013), macromolecular signaling cascades and immunology, economics (Spence, 1973) and financial systems, and cyber security and internet governance (Casey et al., 2019), which in our opinion bodes well that the framework may fit the bill to meet the needs of our cross-scale challenge to understand the neurobiology of memory in particular, and maybe even brain function in general (Rosser, 2003).

In the original (non-cooperative) formulation of game theory (Nash, 1950), the information was considered symmetric between players, in that both players reveal exactly the same information, and use that information in simultaneous strategic choices. Accordingly, in an information symmetric system, information is revealed faithfully, but in more general signaling systems modeled as signaling games, signals can be enhanced with additional or more reliable information. The signal can also be disinformation — deceptive and distorted with respect to the receiver’s expectations (see Fig. 1). One player must therefore respond strategically to the other’s signal, which can keep unrevealed information private. These additional considerations introduce information asymmetry between players, and this asymmetry facilitates more complex interactions between agents.

**Figure 1.**
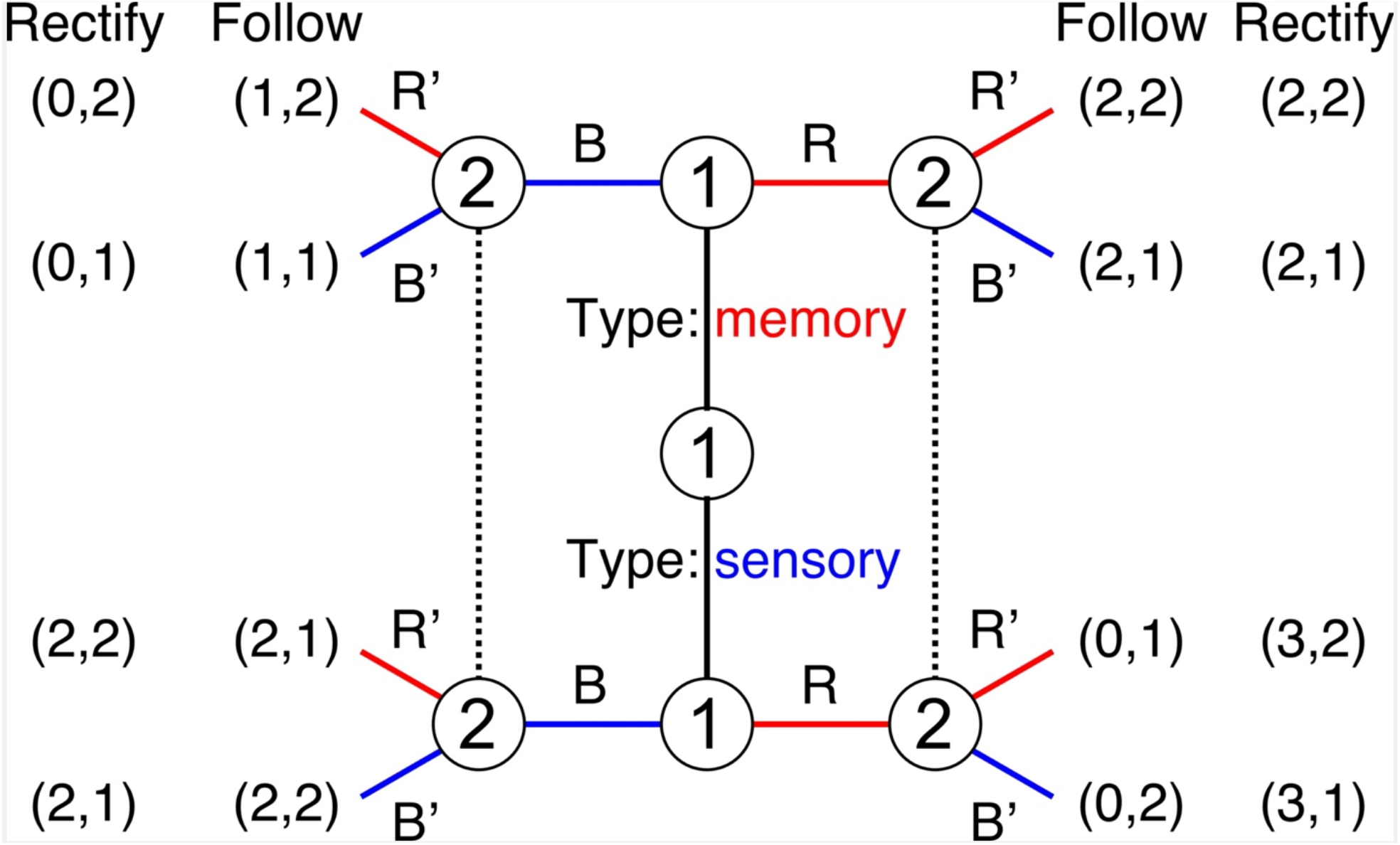
Example. Information asymmetric signaling games illustrate their operation and ability to do information processing based on the payout structure. In this extensive form schematic of a signaling game, the players are two neuronal processing nodes. The sender (player 1) can be one of two types, a memory representation (red) or a sensory representation (blue), which is known by player 1 but not by player 2, which is a motor representation. Player 2 receives an activity signal from player 1 but does not know if it is the sensory or memory signal. The signal can be either pattern R or pattern B and upon receiving the signal, player 2 responds by guessing player 1’s type and generating either motor signal R’ or B’. What signal each player is likely to send depends on the utility of their individual actions, which are represented by the payout coordinates (player 1, player 2) for each of the four possible result pairings of player 1’s type and player 2’s guess R,R’; R,B’; B,B’; B,R’}. Given the set of “Follow” payoffs, when player 1 is type memory, it is likely to signal R (payout=2) instead of B (payout =1) in hopes that player 2 will guess R’, which player 2 will be more likely to do (payout=2) instead of B’ (payout=1). When player 1 is type sensory it is likely to signal B (payout=2) instead of R (payout=0) in hopes that player 2 will guess B’, which player 2 is more likely to do (payout=2) instead of R’ (payout=1). In this “follower” game player 2 correctly reports player 1’s type by generating a corresponding signal. This corresponds to a so-called separating equilibrium because the sensory and memory nodes have adopted distinctive signals, allowing the player 2 motor node to distinguish them, no matter which one generates a signal. Adjusting the payout structure can cause different game behavior. For example, given the “Rectify” payouts, when player 1 is type memory, it is likely to signal R (payout=2) instead of B (payout=0) in hopes that player 2 guesses R’, which player 2 will likely do (payout=2) instead of guessing B’ (payout=1). When player 1 is type sensory it is now incentivized by the payouts to signal R (payout=3) instead of signaling B (payout=2). Sensory is said to signal R dishonestly rather than B honestly, in hopes of deceiving player 2 to signal R’, under the belief that player 1 sensory is actually memory, which player 2 is likely to do (expecting payout=2) instead of guessing B’ (expecting payout=1). Player 2 could have done better if it was not deceived (payout=2), so given player 2 does not know player 1’s type, in this “R-Rectifier” game player 2 will reliably report R’ in response to player 1’s signals, whether player 1’s activity originates from the sensory or memory node. This corresponds to a so-called pooling equilibrium because the sensory and memory nodes have adopted the same signals, preventing the player 2 motor node from distinguishing them.

The quintessential signaling game consists of a sender (informed), a signal (which to varying extents can be informative, uninformative, or misinformative), and a receiver (uniformed; Fig. 1). The receiver is uninformed because it does not directly observe the sender’s type, and so does not know what information the sender knows, all it receives is a signal. As such, the sender type may or may not be correlated with the signal and receiving a signal can change how receivers discriminate between the possible types of sender. These patterns of discrimination by receivers may be considered as “beliefs” regarding what type of sender might be affiliated with a given signal, and since interests are not necessarily aligned between the information-rich senders and information-poor receivers, these beliefs determine the strategies for signaling behavior and may be updated after each subsequent interaction. This viewpoint has consequences for information transmission and the expected “payoff” for each sender and receiver. This latter point highlights that there is feedback coupling between the sender and receiver, which is inherent in a signaling game and observed in neuronal systems (Fig. 2). Note that a receiver in one communication can then act as a sender to transmit signals further through the system in a chain or network-like fashion at each stage, with direct or indirect feedback coupling.

**Figure 2.**
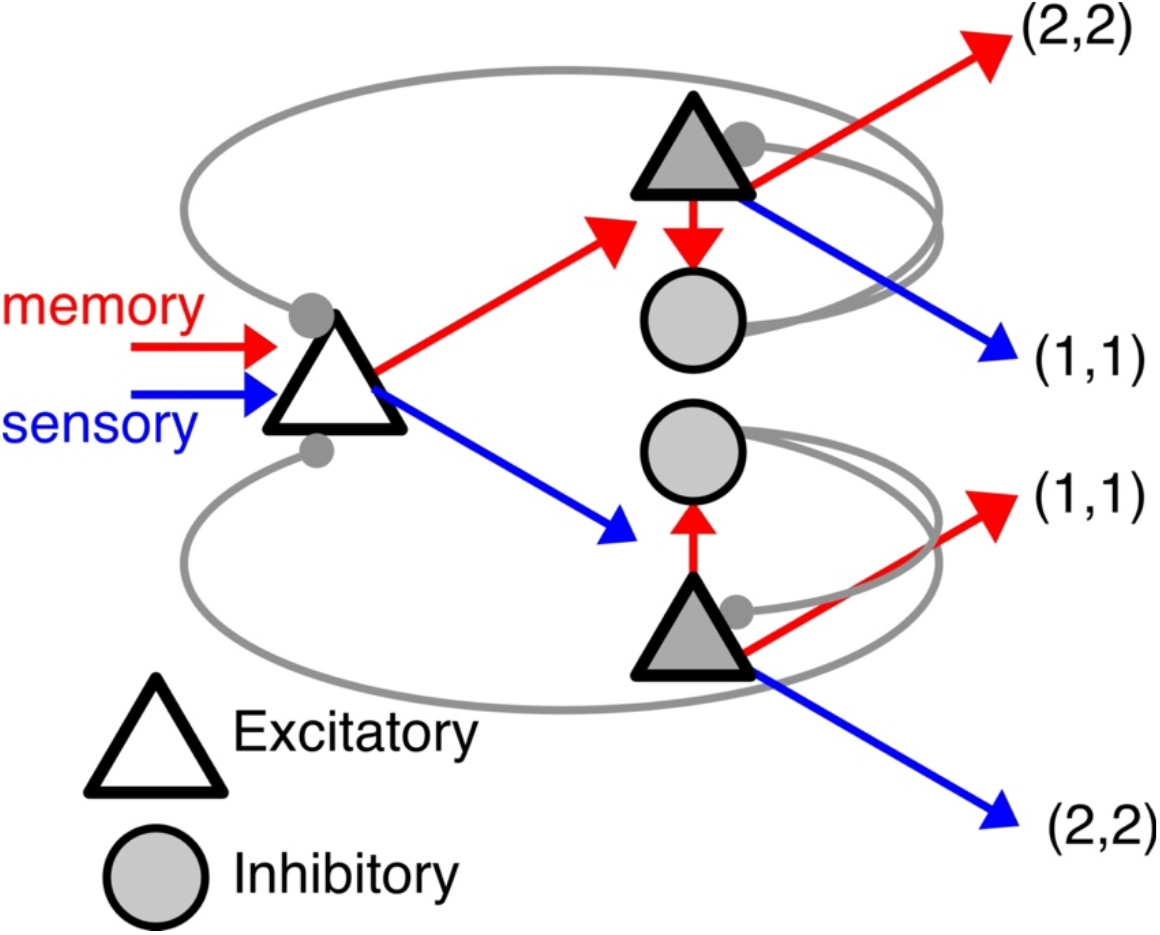
Schematic neural network, where each neural element is a processing node (triangle, circle) and can represent a single neuron or an ensemble of neurons. Given its recent inputs player 1 (open triangle) can generate a representation of type sensory or memory and signal either R or B to the next processing node, the “receiver.” The receiver node is illustrated with feedback and feedforward inhibition via tunable inhibitory synapse-like connections, and it will itself signal either R’ or B’, according to the utilities of the individual processing nodes. The utilities of the two nodes have been set in this example such that the utility of the R and B responses are equal. Consequently, the network will not settle into a stable pattern of signals unless a bias is introduced in the inputs or the connection strengths. The rationale for setting the utility functions for the neuronal network nodes is crucial to the implementation of the game theoretic approach that we are contemplating.

## SIGNALING GAMES

The adaptive utility of relaying a particular signal, in its nascent or modified forms, determines the structure and function of any biological signaling system. In a signaling game, the collection of senders and receivers interact strategically using signals, each trying to maximize their utility. Although the specific instantiations of signaling systems differ, there are general properties of signaling systems that are universal across scales. Fundamentally, the aim of any signaling system is the establishment of conventions. Notice that, in an important way, establishing a convention is different from transmitting information. We highlight this distinction because a standard way of conceptualizing neuronal systems and the brain is as an information processing organ, whereas as we will explain, we are exploring the merits of conceptualizing neuronal systems as signaling organs engaged in signaling games where faithful information processing is just one of many possible states of the system (Fig. 2).

What generates different signaling systems is the information processing that occurs enroute to a particular receiver. In information asymmetric signaling games the sender may know the meaning of the signal, but the receiver is uncertain and possibly completely ignorant about the meaning (Smith, 2000;Rosser, 2003;Dongen, 2006;Jee et al., 2013). As we will discuss presently, this situation represents precisely the signaling condition of the neurons that constitute a nervous system (Fig. 2). In a signaling game, the collection of senders and receivers interact strategically using signals, each trying to maximize their utility. The key to the formalism is a tractable definition of the utility function which determines the behavior of the individual signaling agents. Given each agent in the game has a utility function that it will aim to optimize through signaling, the system of agents is likely to settle into one of several homeostatic states, which in game theory correspond to Nash equilibrium states (Binmore, 2007). A Nash Equilibrium is a profile of strategies such that each player’s strategy is an optimal response to the other players’ strategies. An equilibrium state is explicitly defined as one in which each agent’s behavior is an optimal response with respect to their utility function such that on the whole, no agent can gain more utility by changing their behavior.

It is important to highlight that Nash equilibria correspond to signaling conventions that are adopted by the signaling system, and that these equilibria may or may not promote optimal information transmission across the system. Signaling conventions operate like how social conventions govern the interactions between strangers that meet. To make this essential distinction clear in the context of information processing, consider the notion of an object in a computer program, or a piece of software like a web browser. In each case the software establishes a set of conventions as regards data types and even what operations can be performed on the data types. As in a signaling game, those conventions do not specify or define what specific information is represented or signaled to the receiving piece of code or the user.As in the signaling game in Fig. 1, and as we hypothesize in neuronal systems, those conventions allow for information processing to occur, and constrains what and how information can be processed, because of the conventions, but the conventions do not specify the information itself, that is another matter.

Batesian mimicry in snakes (Casey et al., 2020) illustrates a biological game theoretic example in which conventionally informative signals, meant to signal “possession of dangerous venom,” can be co-opted by snakes that possess the conventionally informative color patterns without the metabolically costly venom that reinforces the convention. The mimics are “deceptive” signaling agents that obtain utility at the expense of the reliable convention. However, the utility is not absolute; it depends on prevalence, the proportion of deceptive agents in the population. A high prevalence of non-venomous types (deceivers) within a population (senders) with conventionally venomous-signaling patterns allow predators (receivers) to adaptively reduce the significance of these color patterns and they consequently increase predation of the snake population. Because increasing prevalence of deception reduces the salience of the signaling convention, and the cost of increased predation is dear, the tradeoff sets an equilibrium limit on the proportion of deceptive agents in a population that is operating under unchanging and stable conventions.

Honest, costly, and deceptive signaling may seem intuitive in the domain of animal evolution, nonetheless, these principles are broadly generalizable, and we argue, they are translatable to the domain of neurobiology. Most neuroscience studies involving signaling games address cognition, honesty, and deception at the level of human social interactions (Jenkins et al., 2016), but the concepts of signaling games have also been applied to analyze biological phenomena at the non-behavioral sub-cellular scale of genes, RNA, and proteins (Massey and Mishra, 2018). Our fundamental conjecture, the central hypothesis of our program is that once the persistence of a signaling convention confers reliable adaptive utility to an organism, then there will be selective pressure against destabilizing the convention. If this is true at the level of social behavior, we surmise it is also true at all levels of biological organization, including the levels of genes and biochemistry, cell biology, development, neuronal circuits, anatomically- and functionally-defined neuronal systems. While the particular signaling games, their utility/cost functions and conventions may differ for each level of analysis, the analytical framework is universal.

We believe the ambition of a universal analytical framework to be an essential and noble goal because, as we indicated at the start, it remains poorly understood how local biochemical interactions give rise to the informative computations that motivate considerations of the brain-computer analogy. We are heartened because, in simple computational models programmed as signaling games involving categorical discrimination of visual inputs, the information used by agents for categorical discrimination of images can be unrelated to the conceptual features that researchers use to define the category (Bouchacourt and Baroni, 2018). Put another way, if naive signaling agents can find mutually beneficial conventions to communicate distinctions, then they can interact locally to optimize their utilities regardless of the nature of those distinctions.

This process of generating conventions through distinctions can reinforce (or destabilize) the convention and have consequences which feed back to the local signaling agent’s toolkit of utility/cost functions. In this manner, the reliability of a convention for transmitting faithful distinctions is what differentiates the types of Nash equilibria found in any given signaling game.

One type of Nash equilibrium is called “babbling,” during which the agents interact via information-poor signals. Such an equilibrium, while not effectively purposeful, allows the system to explore possibilities. By chance, some of these possibilities allow the investments of costly signaling to improve the reliability of signaling, allowing receivers to more reliably predict sender type, raising the utility of some agents, and changing the type of equilibrium to what is called a “separating” equilibrium. Separating equilibria are characterized by stable conventions wherein strategically optimal signaling is distinct for distinct information types (Crawford and Sobel, 1982). In essence, once the agents establish verifiable signals with net utility outweighing the cost of such signaling, then the system will settle into separating equilibria (Sobel, 2007).

Although separating equilibria arise from information asymmetric considerations, they can approximate solutions of information symmetric games too. This is because compatible interests between senders and receivers result in aggregated information by the signaling system, and that itself can confer a selective advantage to the system as a whole. This would be the case for an organism that possesses neurons that self-organize into reliable cofiring assemblies, likely at the substantial expense of “costly signaling,” as we will elaborate presently. In terms of neuronal activity dynamics, it is easy to describe the activity pattern of a competitive network as a separating equilibrium when a subset of cells is vigorously coactive and a competing set of cells have transitioned to being relatively inactive as is observed in attentional and other functional descriptions of cortical networks (de Almeida et al., 2009).Figure 3 illustrates such a competitive network, which by balancing excitation and inhibition has a strong tendency to adopt activity patterns that correspond to separating equilibria (Kaski and Kohonen, 1994).

**Figure 3.**
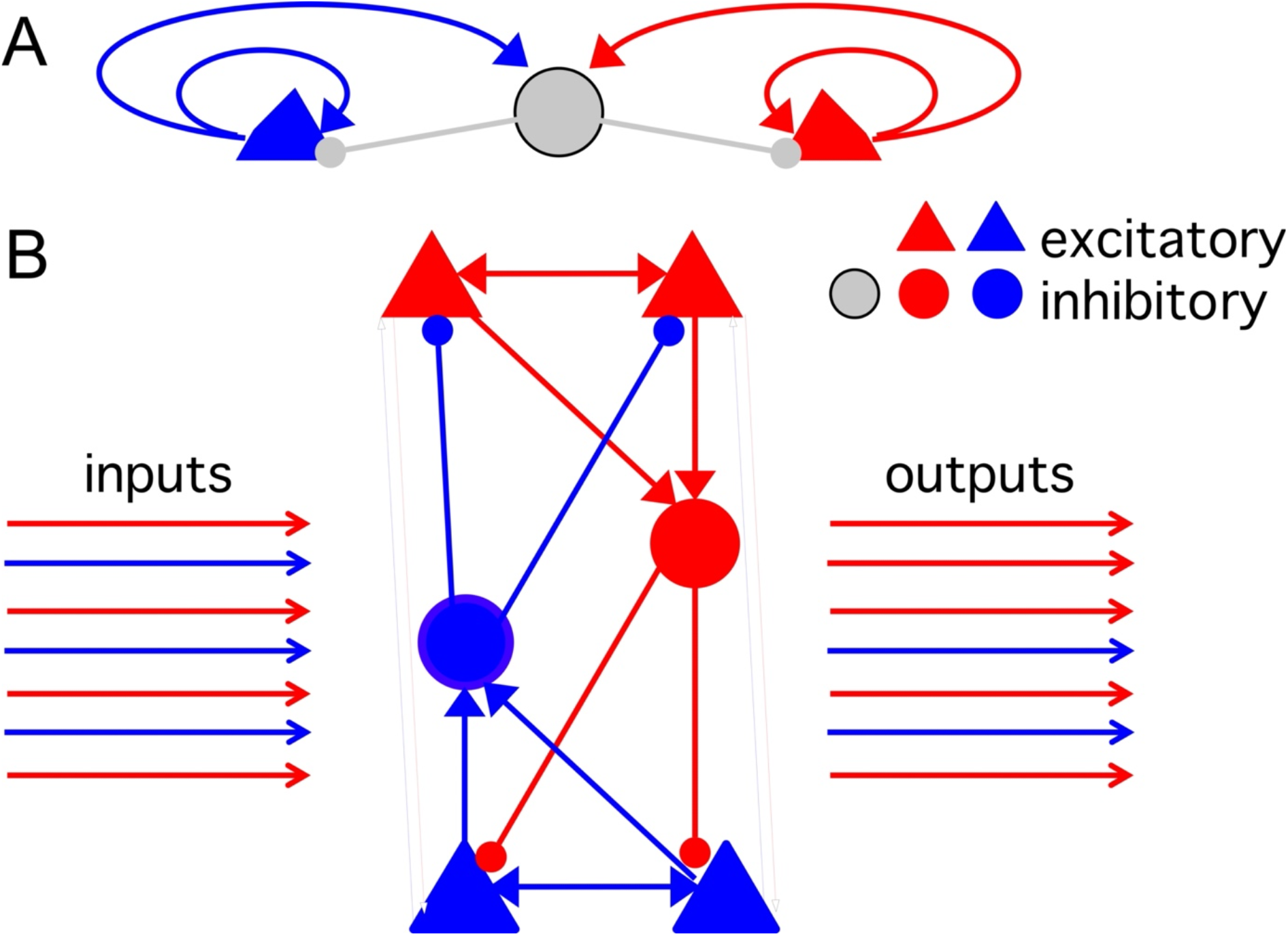
Winner-take-all competitive networks function to select the strongest response amongst a number of competing responses. A) Many network architectures have winner-take-all properties, but the most basic motif is to have sufficient mutual excitation amongst the same-function network elements and mutually-inhibitory coupling between the excitatory elements. B) Illustration of a competitive network where each node represents a population of the nodes. Such a system will transform a set of inputs of varied type into an output that is dominated by type of the strongest inputs.

## INFORMATION ASYMMETRIC SIGNALING GAMES

Information asymmetric signaling game theory assumes no central or overall governance mechanisms, rather that the rules by which the agents behave apply locally and govern local interactions between scale appropriate signaling agents. These conventions are the kinds of local rules like the three that explain much of the behavior and dynamical structures of starling murmurations that can be accounted for by each starling following three rules: 1) keep flying, 2) avoid collisions, and 3) do what the immediate neighbors are doing (Reynolds, 1987;Bialek et al., 2012;Hemelrijk and Hildenbrandt, 2012;Hemelrijk and Hildenbrandt, 2015).

These game theoretic agents can be interacting macromolecules, or pre- and postsynaptic neurons, or neuronal assemblies defined by cofiring neuronal population dynamics. Although we do not elaborate because this work is ongoing, we consider that the utilities that govern the behavior of each neuronal game theoretic agent are bioenergetic functions (Laughlin et al., 1998;Niven et al., 2007). In each case, the system of local, scale-appropriate interactions respects the bioenergetic equilibria that govern their activity. Put another way, dynamical structures (analogous to murmuration configurations) will tend to persist when they correspond to viable signaling conventions. Persistent conventions tend to maintain because like most precedents, they constrain the potential subsequent interactions between agents. This pattern of behavior emerges because most starlings, or biomolecules or neuronal action potential discharge patterns will fail to persistently interact in a manner that is <u>incompatible</u> with the currently persistent conventions — it will simply be too bioenergetically costly to persistently adopt contrarian activities. One can observe this directly in the distribution of cofiring relationships within a population of neurons. The distribution is skewed such that there are many more strongly positive cofiring relationships than anti-cofiring relationships (see Fig. 4). That is however, not to say the anti-cofiring relationships are unimportant. On the contrary, they tend to be rare, their prevalence increases with learning, and they tend to be the most important contributors to the ability to discriminate information from population network activity (Levy et al., 2021).

**Figure 4.**
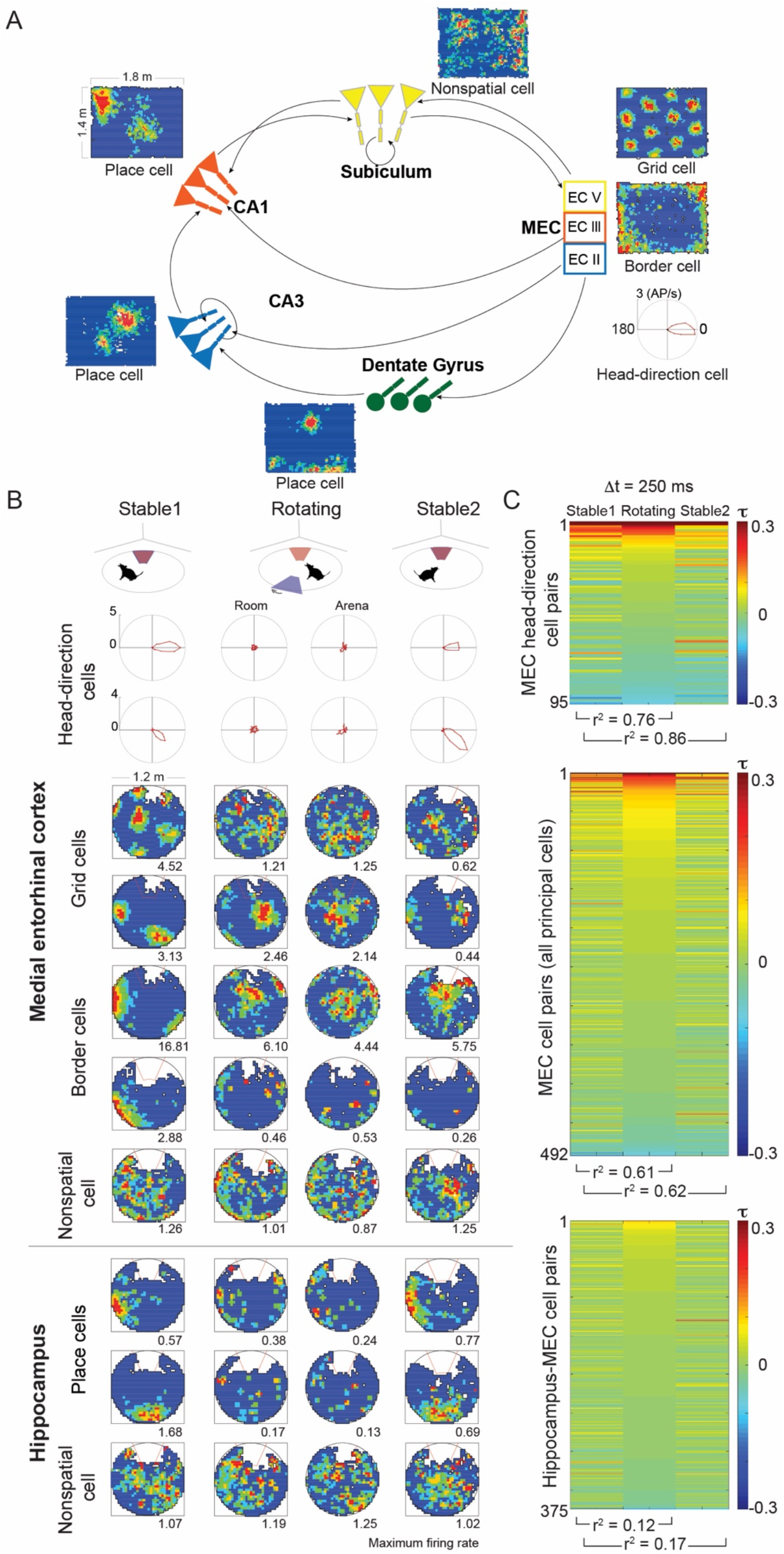
Hippocampal network functional architecture for spatial cognition, and functional cellularization. A) schematic illustrating the functional connectivity amongst the distinct subregions of the medial entorhinal cortex and the hippocampus. Only excitatory inputs are illustrated, emphasizing the interareal connections, whereas the inhibitory connections tend to be mostly local (intra-areal) and have been omitted. Note that the innervations target distinct dendritic compartments. The spatial firing properties of principal cells in each area are also indicated by an example session-averaged firing rate map from neuronal recordings while a rat explored a rectangular arena. Note how grid, head direction, and border spatially-tuned cells of MEC represent component variables from which place can be computed (distance, azimuthal direction, and environmental boundary, respectively) from the spatially-tuned MEC inputs to the subfields in which place cell spatial firing patterns are the most frequently observed spatial firing patterns. B) Top-to-bottom: Schematic of stable-rotating-stable experimental active place avoidance experimental conditions. Example polar firing rate representations of two head-direction cells and blue-to-red color-coded firing rate maps illustrate the typical spatial tuning of MEC and hippocampus (CA1) cells recorded while rats navigate during two-frame place avoidance in a stable-rotating-stable triad of 30-min recordings. The rotating session dissociates the accessible space into two spatial frames, a stationary room frame and a rotating arena frame, and the two frame-specific firing rate maps are provided for each cell. The number under each map is the minimum rate in the red category measured in AP/s units. Note how the spatial tuning observed during the stable sessions is lost during the rotation. C) Distribution of cell pair cofiring measured as Kendall’s correlation (τ) computed at 250 ms resolution. Each simultaneously-recorded pair of cells was recorded in the stable-rotating-stable triad session. The cell pairs are sorted in descending order by the value of τduring the rotating recording and the order of cell pairs is maintained for all three recordings. Note (1) that the correlation patterns skew to significant positive values with few significant negative values, and (2) that the correlation patterns strongly persist across the session triad (r^2^ coefficient of determination values given below the plots) for MEC recordings. In contrast, the persistence of cofiring is much weaker for MEC-hippocampus cell pairs (Stable – Stable: MEC-MEC pairs (r = 0.79) vs. MEC-HPC pairs (r = 0.41) z = 9.24, p ~0; Stable-Rotating: MEC-MEC pairs (r = 0.78) vs. MEC-HPC pairs (r = 0.35) z = 9.88, p ~ 0). See Appendix 1 for Methods.

Once established, persistent conventions tend to further increase their persistence. Consider those agents that are currently behaving in ways that are independent, or even contrary to a currently instantiated convention, for example the neurons with weak synaptic connections that, as a result, discharge independently of the dominant neuronal network pattern. Such neurons may be subsequently recruited to the dominant network discharge pattern. Put another way, they may adopt a discharge pattern that mimics the dominant discharge pattern and then through the costly signaling of synaptic plasticity, become legitimately incorporated into the persistently active network pattern of discharge. One might expect in this case that the weakly cofiring neuron pairs will increase their likelihood to cofire. Although such recruiting of independent neurons into the persistent activity of an established cell assembly appears intuitive, perhaps even routine, an interesting, surprising consequence of this recruitment can occur when the signals of the recruited neurons also provide information that is distinctive from the information that the network processes with its signaling convention. This is in fact observed in the cofiring relationships of hippocampus principal cells under acute intoxication with the psychotomimetic phencyclidine (Kao et al., 2017), or other cognition impairing manipulations, such as TTX-induced disinhibition of the contralateral hippocampus (Olypher et al., 2006). In both cases, cell pairs with negative or weak cofiring statistics before the drug manipulation, increase their cofiring under the manipulation, whereas the initially strongly cofiring cell pairs are unchanged. This pattern of selectively increased cofiring coincides with the inability to perform behavioral tasks that require discriminating between relevant and irrelevant information, but the same cofiring increase is not impairing in conditions where the irrelevant environmental information is attenuated (Wesierska et al., 2005;Olypher et al., 2006;Kao et al., 2017). From the perspective of the information that that the neuronal activity represents, by adopting the convention, the recruited neurons could have deceived the neuronal network partners that they have been recruited to join. This framework has interesting, and powerful explanatory implications, we will discuss shortly. For the time being, it provides an explanation of observed increase in cofiring of initially anti- and weakly cofiring cell pairs after knowledge impairing manipulations. This game theoretic perspective can also explain another otherwise puzzling observation from ensemble recordings of spatially-tuned neurons in the entorhinal cortex and hippocampal regions that are thought to constitute the brain’s navigation system (Fig. 4).

## ENTORHINAL-HIPPOCAMPAL NEURONAL POPULATION DYNAMICS: PHENOMENA IN NEED OF A CONCEPT

We will now elaborate on the entorhinal-hippocampal neuronal system to set the foundation for what we aspire to understand and explain using the concepts of information asymmetric signaling games (Fig. 4A). We focus on hippocampus neuronal population discharge correlates of spatial information that can be measured by studying the spatial behavior of freely-behaving rodents, as this is our long-standing experimental research program (see Methodology in Appendix 1). In freely-behaving rodent subjects like rats and mice, an environment-specific 20-25% subset of hippocampus principal cells discharge action potentials robustly only when the subject is in cell-specific locations called the cell’s firing field (O’Keefe, 1976); place cells are also identified in birds and bats (Ulanovsky and Moss, 2007;Yartsev Michael and Ulanovsky, 2013;Payne et al., 2021). When neurons discharge in this way they are traditionally called “place cells” (Fig. 4A,B). The entorhinal cortex contains neurons that signal distance travelled (“grid cells”), head-direction (“head-direction cells”) (Sargolini et al., 2006), the presence of environmental borders (“border cells”) (Solstad et al., 2008) and the current speed of locomotion (“speed cells”) (Kropff et al., 2015). The 2014 Nobel Prize was awarded for the discoveries of these functional cell classes (Fenton, 2015;Moser et al., 2017). These spatially-tuned MEC neurons project to the hippocampus in multiple, parallel and distinctive pathways, providing multiple sources of the information components for computing “place” (Baks-Te Bulte et al., 2005;Witter, 2006;2007;Canto et al., 2008)(Fig. 4A).

We developed an experimental paradigm in which rats and mice readily navigate on a slowly, continuously rotating circular arena (Fig. 4B top). The arena rotation dissociates the accessible space into two simultaneous and distinct spatial frameworks. One is stationary defined by room-anchored landmarks, and the other is rotating, defined by arena-anchored stimuli such as scent marks on the rotating surfaces. Animals quickly demonstrate that they understand the room and arena spaces to be distinct, even if the arena never rotates (Fenton et al., 1998;Fenton and Bures, 2003). They demonstrate this knowledge in the active place avoidance paradigm, in response to training during which we punish the animal with a mild electric shock for entering a room-defined zone and/or an arena-defined zone (Fig. 4B top; Fenton et al., 1998;Fenton and Bures, 2003). In response, they will quickly and selectively avoid the room and/or arena shock zone, which is why we call the behavior two-frame place avoidance. Two-frame place avoidance is one of the most sensitive tasks to disturbed hippocampal function; even inactivating one of the two hippocampi makes the animals unable to learn, consolidate, or remember the location of the shock zone (Cimadevilla et al., 2000;Cimadevilla et al., 2001;Kubik and Fenton, 2005;Wesierska et al., 2005;Kelemen and Fenton, 2010).

We investigated whether during two-frame place avoidance, the spatially-tuned cells in hippocampus and entorhinal cortex would exhibit spatial tuning in the room frame or the arena frame. Despite the animal navigating the rotating environment extremely well, very few neurons (a fraction of a percent) demonstrate their place cell, grid cell, or head-direction cell spatial-tuning properties during rotation; the tuning returns once the rotation stops (Fig. 4B). We have shown that this apparent loss of spatial tuning is because the neuronal populations collectively signal the animal’s location and direction in an internally-organized, multistable manner such that the ensemble activity patterns switch between representing the current location and direction in <u>either</u> the room frame or the arena frame, but not both (Kelemen and Fenton, 2010;Kelemen and Fenton, 2013;Talbot et al., 2018;van Dijk and Fenton, 2018;Park et al., 2019;Chung et al., 2021). Multistable switching between room and arena representations is rapid (sub-second) and occurs in a purposeful way with a periodicity in the range of 10 seconds (Kelemen and Fenton, 2010;Kelemen and Fenton, 2013;van Dijk and Fenton, 2018;Park et al., 2019).

These multistable population discharge dynamics in maintained environmental conditions indicate that neuronal activity is internally-organized and variably registered to the environment in a manner that takes into account the distinctive spatial frames and registers neuronal ensemble activity to the spatial frame that is more currently useful (i.e., use the room frame when near the stationary room-frame shock zone and use the arena frame when near the rotating, arena shock zone). The cofiring relationships of simultaneously recorded cell pairs provides additional, independent evidence for this strong internal organization of neuronal discharge patterns. This conclusion can be arrived at by computing the pairwise correlations among all simultaneously recorded cell pairs (Schneidman et al., 2006). During stable and rotating recordings, we observe that the pairwise coactivity measures of their correspondence is indistinguishable. An example from the head-direction cells recorded from superficial layers of the medial entorhinal cortex is shown in Fig. 4C top, indicating that the population discharge of these “same-function” neurons is dynamically rigid (stationary and steady) (Park et al., 2019). A similarly persistent set of cofiring relationships is also observed for all the simultaneously recorded neurons in superficial medial entorhinal cortex (Fig. 4C middle), further indicating that despite mixed tuning to different spatial variables (Park et al., 2019), as a whole, the MEC network manifests the stationary and steady temporal discharge dynamics that are characteristic of neuronal attractor dynamics (Yoon et al., 2013;Chaudhuri et al., 2019). Like MEC, the population discharge properties of hippocampus also exhibit attractor dynamic properties, in that the collective activity of MEC cells, or the collective activity of the cells of the hippocampus CA3 or CA1 subfields tend to exist as relatively stable patterns of activity that are readily described as a low-dimensional manifold in the high-dimensional activity space that is defined by the independent activities of the population of cells (Samsonovich and McNaughton, 1997;McNaughton et al., 2006;Yoon et al., 2013;Chaudhuri et al., 2019;Gardner et al., 2021;Nieh et al., 2021). Evidence of these manifolds of low-dimensional neuronal activity can be readily measured as conserved pairwise coactivity patterns across the network, which are known to represent higher-order correlations (Schneidman et al., 2006;Levy et al., 2021) (Fig. 4C).

## DO SIGNALING GAMES OFFER ANYTHING THAT INFORMATION MAXIMIZATION DOESN”T ALREADY?

Our observations of multistable frame-specific positional and directional population discharge, and persistence of internally-organized cofiring discharge relationships (Kelemen and Fenton, 2010;Talbot et al., 2018;van Dijk and Fenton, 2018;Park et al., 2019) can be readily accommodated by the information maximization (Infomax) perspective that is a dominant conceptualization of how information processing is serially organized across connected neuronal networks with related input-output functions (Bell and Sejnowski, 1997;Linsker, 1997), such as the MEC and hippocampal computations of environmental space (Solstad et al., 2006).

Neuronal ensemble discharge patterns are assumed to be the neuronal representations and or instantiations of mental objects, in short, the expression of cognitive phenotypes. This view assumes that generating reliable representations of cognitive variables essentially constitutes proper entorhinal-hippocampal ensemble function. Such assertions emerge from the Infomax hypothesis that has been important for systems neuroscience because it offers a unifying principle (Barlow, 1961). Accordingly, neuronal computations at each level of the nervous system would operate to maximize the mutual information between inputs and outputs that the computation is operating on, and so couple the information content across a series of such neuronal computations, the output of one serving the input to the next. While Infomax principles are readily applied to predict what neuronal discharge might signal, it is difficult to apply these principles to the biochemistry and cell biology that operates and underlies the neuronal discharge that define the mental objects of interest. Furthermore, Infomax concepts are severely challenged to explain our additional observation that the cofiring relationships between simultaneously-recorded hippocampal and MEC neurons (Fig. 4C bottom) is only weakly persistent across the stable and rotating arena conditions, despite both the entorhinal (Park et al., 2019) and the hippocampal populations each exhibiting strongly persistent, attractor-like cofiring relationships (Levy et al., 2021). If MEC representations of space project to hippocampus then their discharge should be informationally coupled, and the discharge representations should be informationally coherent, perhaps even increasing in fidelity according to Infomax predictions. But as Fig. 4C suggests, the discharge coupling is unstable, and the frame-specific multistability in hippocampus (Kelemen and Fenton, 2010;van Dijk and Fenton, 2018) is observed to have a different, weakly opposite relationship to frame-specific behavior as is observed for MEC frame-specific multistability (Park et al., 2019). Because it is difficult to reconcile these observations within the Infomax framework, we were motivated to seek an alternative. We contend that the framework of signaling games has utility for understanding these challenging features of hippocampal neuronal population dynamics at the level of systems neuroscience, as well as at other levels of biological organization, including cell biology and biochemical signaling that others have demonstrated (Smith, 2000;Jee et al., 2013).

The game theoretic framework of fundamentally coupled senders and receivers provides a natural analytical language and framework for characterizing these neuronal dynamics and for analyzing their multiple steady states as game theoretic Nash equilibria. Indeed, the manifold patterns of attractor-like neuronal activity (Amit, 1992) are readily described by the concept of “cellularization” that naturally emerges in the game theoretic language. Through reliable and oftentimes costly signaling like metabolically-instantiated synaptic plasticity, multiple game theoretic agents, in this case neurons within an interconnected network, can settle into stable population equilibrium states that are characterized by signaling conventions within the network acting as a single collective. In other words, robust neuronal population dynamics produce activity patterns that themselves act as game theoretic senders and/or receivers, adopting the form of a dynamically “cellularized” object, despite being composed of individual neuron components. Note that this process of cellularization naturally defines cross-scale interactions and information transfer that are seamlessly transcended by the game theoretic formalism, whereas such cross-scale analysis have been a challenge for other frameworks and conceptualizations.

## ENTORHINAL-HIPPOCAMPAL NEUROBIOLOGY AS AN INFORMATION ASYMMETRIC SIGNALING GAME

We believe that the ideas of information asymmetric signaling games can readily and advantageously apply to neuronal systems. To that end we now reconceptualize the neurobiological interactions that were described above in these game theoretic terms. Simple conserved game conventions can first arise from arbitrary dependencies, much like how spuriously causally coactive neurons can undergo synaptic plasticity so they are more likely to cofire in the future, and in so doing establish a neuronal firing sequence convention. Selection for useful conventions can then privilege those with utility so they persist with potentially increasing complexity, much like standard reinforcement learning models and Hebbian learning rules assert.

It may be difficult to appreciate how multi-stable cognitive organization can emerge from agents interacting locally in simple games. It is therefore important to emphasize how slight deviations in initial conditions can create lasting differences in evolving dynamical systems. For instance, in area CA1 of the hippocampus, multiple representations of the same spatial environment can coexist as recurring stable patterns of firing (Sheintuch et al., 2020). However, with time and additional exposure to the environment, these representations tend to deviate, according to the signaling game framework, because the information asymmetry among the representations generates persistent distinctions.

Mechanisms to persist, to maintain persistence and to increase complexity of signaling conventions typically require metabolic and other resources and so come at a cost that most agents will not engage (Niven et al., 2007). Such exclusions of non-compatible agents increases the reliability of the signaling system as a whole, which can make costly signaling advantageous (de Ruyter van Steveninck and Laughlin, 1996). Synaptic plasticity is metabolically costly, and molecular biosynthesis needs to be coordinated pre- and postsynaptically for the structural plasticity that changes the shapes and sizes of synapses engaged during learning. This coordination results in a myriad of local processes including actin polymerization, synthesis and then translocation of synaptic proteins. In addition, once strengthened, potentiated synapses will result in greater transmembrane ion currents and thus dissipation of the sodium and potassium ionic concentration gradients, which, to be restored, will require increased Na/K ATPase biosynthesis and activity. Indeed, the upregulation of PKMζ, which is necessary and sufficient for the long-term maintenance of wildtype LTP at hippocampal and other synapses, coincides with the upregulation of the Na/K ATPase (Tian et al., 2008). This observation is consistent with the notion that the increased communication comes at an energetic cost and so once in place, is likely to contribute to the information asymmetry.

As mentioned previously, information symmetric games are those in which both players reveal complete information about each other and consequently are confident about how the other rational players will strategically respond in each scenario. Information asymmetry occurs when one player (the sender) possesses information about its signaling type that is not available to the other player (the receiver). By introducing information asymmetry, we can better describe molecular signaling cascades and neuronal networks. As an example of a cascade, consider the multiple roles of intracellular calcium, which can initiate postsynaptic vesicle release, or changes in gene expression, or programmed cell death, each depending on the origin of the calcium. In the case of a neuronal network, considering that each neuron receives ~10^4^ distinct synaptic inputs, the consequences of their activation are integrated to change the transmembrane potential at the remote location of the axon hillock. The neuron’s axon hillock is uncertain about which subset of the ~10^4^ synapses activated in temporal and spatial summation, causing it to generate an action potential. This fundamental uncertainty creates information asymmetry and the opportunity for game theoretic deception.

Deception occurs when there is information asymmetry and insufficient common interest between the sender and receiver. Costly signaling may be exploited by deceptive agents exploiting the convention, even though they degrade the reliability of signaling. Examples of these preserved conventions include canonical neurotransmitters and receptors, ubiquitous membrane proteins, and immediate early genes. Examples of deceptive agents include but are not limited to pharmacological or natural agents that mimic the biomolecular structure of these canonical agents. These conventions are ubiquitous across vertebrates and can be found functioning in both *in vitro* slice preparations as well as in *in vivo* preparations such as recordings from awake freely-behaving subjects. Essentially the same local cell-biological biochemical constraints underlie the structural dynamics of all neuronal populations during transcription, translation, membrane depolarization, action potential generation, and synaptic and neuronal population synchrony that is measured as oscillations in local field potentials and dynamic cofiring patterns in neuronal ensemble activity. While not exhaustive, we believe these examples illustrate there is substantial and natural potential to translate neurobiological phenomena and theory into the formal and descriptive language of information asymmetric signaling game theory. In fact, as we alluded to above, a game theoretic framework offers mechanistic accounts for neuronal network phenomena that are otherwise difficult to explain, as we will now illustrate, by applying the concept of deception.

## DECEPTION: INFORMATION INCOMPLETENESS VS INFOMAX

Let’s consider the notion of game theoretic deception in the context of an explicit neuronal discharge phenomenon that we have studied in the neuronal dynamics of the hippocampus. Hippocampal neurons, residing in the CA1 output subfield, can discharge as if to represent the current location of a mouse. This interpretation of the observables leads one to think of the “place cell” phenomenon. From there it is relatively straightforward to understand how a place cell network of neuronal activity is “encoding” the current location. However, we also observe that, at least transiently, the same CA1 neuronal network’s discharge will not signal the current location. Instead for about 500 ms, the discharge will represent a distant location that, during a spatial memory task, can be shown to represent the mouse’s recollection of a remote goal or some other behaviorally-important location (Fig. 5). Conventional explanations do not easily explain how such non-local place cell activity can come about, and the field has hypothesized that there might be specialized “goal” or “reward” cells, rather than place cells that sometimes act like goal or other types of functionally-defined cells (Poucet and Hok, 2017;Gauthier and Tank, 2018;Duvelle et al., 2019). Recent work shows that any place cells can transiently express non-local place cell discharge that otherwise resembles routine locally-legitimate place cell firing. It is as if the network has been deceived into signaling a remote location. The deception appears especially strong and maladaptive in *Fmr1-null* mutant mice that model the genetic defect in human Fragile X Syndrome, the leading cause of intellectual disability and autism. *Fmr1-null* mutant mice express excessive synaptic plasticity in hippocampus CA1 because of memory training (Talbot et al., 2018). When challenged with a novel location after learning an initial location, *Fmr1-null* CA1 place cell network discharge and the mouse’s behavior, both recollect the formerly correct location rather than the currently correct location for the memory task (Dvorak et al., 2018). These phenomena are straightforward to explain as instances of deception within asymmetric signaling games, and the neuronal mechanism for transiently switching the hippocampus information processing from encoding to recollection is consistent with this game theoretic explanation (Dvorak et al., 2021). There is a transient, strong synchronous discharge event that originates in an upstream region, the medial entorhinal cortex (MEC) and is observed in the dentate gyrus (DG). The event is called a MEC-originating dentate spike (DS_M_). The DS_M_ acts like a control switch for the MEC → DG →CA3 → CA1 neuronal circuit (Fig. 5) that changes information processing in the circuit in a way that causes the CA3 cells to discharge in a manner that is informationally disconnected from the ongoing location-specific discharge in the dentate gyrus. The DS_M_-triggered CA3 activity appears to deceive CA1 into discharging in a manner that appears normal, only, instead of representing the current location, the DS_M_-associated CA1 discharge represents a remote, recollected location (Fig. 5). A similar, non-local place cell ensemble discharge phenomenon has also been described that is easily explained by this deception concept. Again, on a subsecond timescale, the ensemble discharge of CA1 neurons will toggle between representing non-local places that a rat may visit in the near future (Kay et al., 2020). It is as if the activity during that moment represents something akin to the rat’s aspirations or intentions rather than where it is, which is not easy to explain from a feed-forward information maximization conceptualization of hippocampal spatial information processing but is a natural consequence of deceptive signaling.

**Fig 5.**
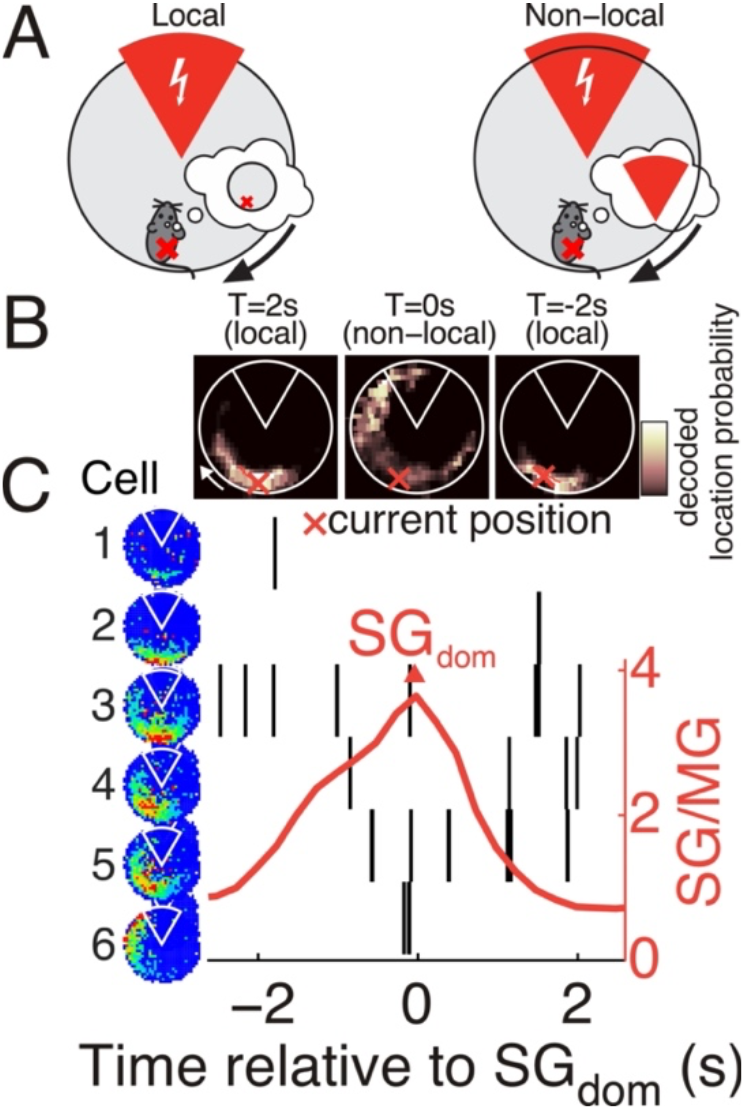
Non-local hippocampal network discharge as deception. A) Cartoon illustrating that hippocampus place cell activity can typically be decoded to accurately identify the subject’s current position (left). However, at times the position decoding is non-local in that it points to a position remote from the subject’s current position (right). This non-local decoding is often to the vicinity of a shock zone (red sector) if the mouse has been trained to avoid that shock zone. Furthermore, the non-local representation of position tends to occur ~2 seconds before the subject will make avoidance movements away from the shock zone, suggesting that it is a recollection of the shock zone’s location. B) Bayesian posterior maps of the likelihood of a mouse’s position calculated on the basis of a 500-ms observation of place cell population discharge centered at the times indicated. C) T=0s is the moment that slow gamma dominance was observed in the concurrent CA1 local field potential (SGdom). SGdom is computed as the ratio of slow (~30 Hz) to mid-frequency (~70 Hz) gamma (MG) in the local field potential, and T=0 is marked as the peak of the sufficiently positive SG/MG ratio. The firing rate maps of six place cells are shown ordered from the one with the firing field closest to the mouse’s current location to the one with the firing field distant from the current location and near the shock zone. Note that during this 5-s data segment, that despite the mouse moving very little, which place cells discharge action potentials (rasters shown) changes from the cells with fields at the current location, to the cells with fields near the shock zone, remote from the mouse. It is not obvious what sensory cues would switch firing from representing local position to non-local position, but SGdom transiently switching CA1 network function from processing local place information based on sensory cues to processing non-local information based on memory predicts these observations. SGdom is triggered by a medial entorhinal cortex-originating dentate spike event that switches hippocampal information processing from sensation-based encoding to memory based recollection (Dvorak et al., 2021). Figure based on (Dvorak et al., 2018).

## WRAPPING UP

We have recognized that signaling game theory can successfully explain, describe, and model complex phenomena in diverse fields ranging from biomolecular evolution to economics. From the small-scale to the large-scale, at each level of analysis there are biological signaling agents transmitting signals in attempts to maximize their respective utility. Information asymmetry between signaling agents can lead to stable equilibria under cooperative circumstances, equilibria which may persist due to the low cost of signaling relative to the acquired utility of signaling. These persisting signaling equilibria are considered conventions. Signaling conventions are often preserved across species, and interspecies analogues add credence to the universality of signaling systems as concrete neurobiological objects of study.

Neurons use biochemical and bioelectric signals to differentially allocate metabolic resources to particular “privileged” connections. Although the local metabolic constraints of neurons are determined by biochemical considerations, natural selection and experience can act on these asymmetries at the level of their emergent features. These emergent features are phenotypes, and for our consideration these are the phenotypes of cognition and behavior, which are correlates of neuronal ensemble discharge patterns.

Coordinating behavior into coherent patterns is a unifying feature of all evolutionary signaling systems, and the reliability of signaling is determined by how well a signal can predict a sender”s type. For instance, a signaling system approximating Infomax consists of senders reliably signaling their type, thereby reducing distortion, and generating optimal and predictable responses. But, as mentioned above, observable, and adaptive hippocampal function demands more than optimally reliable information. Spatial information maximization is instead merely one end in a spectrum of signaling organization: the optimally informative end. On the other end of the spectrum is completely uninformative signaling (so-called babbling equilibria) whereby signals do not reliably distinguish among sender types. Either end of this spectrum alone is untenable for the demands on hippocampal function. Robust signaling conventions cannot reliably emerge through signaling that is uninformative to the overall system, and maximally stable signaling conventions cannot flexibly evolve under an information maximization framework.

It is our contention that entorhinal-hippocampal organization can be understood as the cellularization of scale-specific signaling agents ranging from genes, proteins, neurons, and neuronal populations that interact according to shared signaling conventions. Every cellularized group of signaling agents is organized somewhere along the babbling ↔ separating signaling spectrum. From the scale of neuronal ensemble dynamics, and emergent sensorimotor features, information-rich separating equilibria can approximate information maximization paradigms which have been highly fruitful to systems neuroscience research. These populations of highly cofiring neurons are well suited to encode reliable sensorimotor features of the world, allowing an organism to benefit from the high degree of cellularization and reliable signaling.

Furthermore, signaling game theory also provides an explanation for aspects of hippocampal activity which are not well understood from a systems level information maximization paradigm. Populations of cofiring neurons further from separating equilibria generate less cellularization, allowing signaling conventions to adapt to changing relations between the organism and the world. We observe that loose correlations observed in less cellularized populations underlie *Fmr1-null* mutant CA1 hippocampus’ susceptibility to deception (Talbot et al., 2018). Moreover, the dynamic restructuring of population vector correlations observed during remapping experiments in area CA1 (Redish et al., 2000;Jackson and Redish, 2007;Kubie et al., 2020;Levy et al., 2021) provide further support for the significance of signaling equilibria that can transiently stray from separating equilibria to establish new conventions. In summary, the tension generated between different cellularized agents and their respective level of cellularization can account for the complex array of functions attributed to the hippocampus, functions which emerge from metabolic and energetic constraints of individual neurons and scale up to population-level phenotypes of cognition and memory.

## Acknowledgements

We are grateful to Will Casey, Cristina Savin and Kshaunish Soni, and Mike Wigler for discussions.

## APPENDIX 1: Methods

These procedures were previously described in detail (Park et al., 2019), and are here provided in brief to complement the data reported in Figure 4.

### Subjects

We used 14 adult male Long-Evans hooded rats weighing 300-400 g (Taconics Farms, NY). All experimental procedures were approved by NYU’s Institutional Animal Care and Use Committee. Rats were handled by the experimenter for 5 days (5 min/day) before surgery under pentobarbital (50 mg/kg, i.p.) anesthesia. The surgery was to implant custom microdrives on the skull that could micro-position electrodes at the recording sites in the bran. Two-four independently movable tetrode-configured electrode bundles were targeted to the hippocampus (relative to Bregma AP 3.8, ML -2.5, DV 1.9) and the medial entorhinal cortex (relative to the from sinus AP 0.5, ML, 4.5, DV, 2.0) of seven rats. Eight independently movable tetrode electrodes were targeted to the medial entorhinal cortex of seven rats. The animals were allowed at least a week to recover before behavioral training began.

### Behavior

The rats were habituated to the stainless-steel disk arena (diameter 1.2 m). They were encouraged to continuously forage for 20-mg sugar pellets (Bio-Serv, NJ) that were randomly dispensed to random locations by a computer-controlled overhead feeder. Each of the 5 days of the habituation phase, the arena was stable for 30 min and rotating at 1 rpm for 30 min. Active place avoidance training followed to avoid an annulus-sector that for some rats was 30° and for other rats was 45°. The sector extended from the edge of the arena toward an annulus at either 50% or 40% of the radius, respectively. The rats were trained to avoid a mild <0.4 mA foot shock if they entered a shock zone that was defined in a fixed location of the room and a coincident fixed location on the arena surface. When the arena was stable the room-defined and arena-defined shock zones were physically identical, but when the arena rotated, they were dissociated such that the stationary room location of the shock zone remained fixed in the room (but not on the rotating arena), and the rotating arena location of shock remained fixed on the arena (but not in the stationary room). The rats received ten daily training sessions consisting of the triad stable1-rotating-stable2, each lasting 30 min, where the arena was stable or rotating and the two stable sessions were identical with the area in an identical orientation. Using two infrared LEDs mounted to the recording electronics on the rat’s head, the rat’s position and head direction were tracked 30 times a second using an overhead video camera and software (Tracker, Bio-Signal Group, Acton, MA). Position and head direction were tracked in both the stationary spatial frame of the room and the rotating spatial frame of the arena by referencing an infrared LED that was attached to the arena.

### Electrophysiology

The electrodes were advanced into the CA1 and MEC regions until action potentials could be recorded. The signals were filtered between 300 Hz and 7 kHz, amplified up to 7000 times and digitized at 48 kHz using commercial hardware and software (dacqUSB, Axona Ltd, St Albans, UK). In this way, ensembles of action potentials were recorded from dorsal hippocampus and dorsomedial entorhinal cortex (MEC) while the animals performed the active place avoidance task in the stable and rotating conditions. A total of 1465 single units were recorded from MEC and 756 single units were recorded from CA1. Single unit isolation quality was assessed by computing IsoI_BG_ and IsoI_NN_, the Isolation Information measures (Neymotin et al., 2011). Only the single units that had values greater than 3.5 bits were considered sufficiently well-isolated for these studies (MEC 810, CA1 414).

### Data Analysis

Custom C/C++ and MATLAB software was used to compute all outcome measures, with details published, and code publicly available (Park et al., 2019). Functional cell classes were determined by statistical criteria after computing firing-rate tuning as a function of place and head-direction determined by dividing the number of action potentials that a single unit discharged while the rat was in each 2.13-cm square positional bin or 10° directional bin divided by the total time the rat was detected in that bin. MEC head-direction cells were classified as single units with long duration (> 350 us) action potential and direction-independent firing rate below 10 AP/s. CA1 place cells were classified as single units with long duration action potentials (> 350 us), position-independent firing rate < 5 AP/s, and spatial coherence (> 0.4). The network stability was estimated by computing the n(n-1)/2 pair-wise spike train correlations between all simultaneously recorded pairs of n cells. The spike counts were computed in fixed duration 250 ms time bins for each spike train and Kendall’s correlation (τ) was calculated between the spike counts of all pairs of simultaneously recorded cells within MEC, CA1, and between MEC and HPC. The stability of the network state across the Stable1 vs Stable2 and across the Stable1 vs. Rotating conditions was investigated by comparing the τ values for each cell pair in the two conditions, and this was quantified by computing the Pearson correlation across the pairs of τ. The coefficient of determination (r^2^) is used to estimate the variance in the τ values in one condition that is explained by the variance in the other condition, and thus the stability of the network discharge. Statistical comparisons between distributions of correlation were performed after transforming r values to z using Fisher’s transform.

